# Exploring the spatially explicit predictions of the Maximum Entropy Theory of Ecology

**DOI:** 10.1101/003657

**Authors:** D.J. McGlinn, X. Xiao, J. Kitzes, E.P. White

## Abstract

**Aim:** The Maximum Entropy Theory of Ecology (METE) is a unified theory of biodiversity that attempts to simultaneously predict patterns of species abundance, size, and spatial structure. The spatial predictions of this theory have repeatedly performed well at predicting diversity patterns across scales. However, the theoretical development and evaluation of METE has focused on predicting patterns that ignore inter-site spatial correlations. As a result the theory has not been evaluated using one of the core components of spatial structure. We develop and test a semi-recursive version of METE’s spatially explicit predictions for the distance decay relationship of community similarity and compare METE’s performance to the classic random placement model of completely random species distributions. This provides a better understanding and stronger test of METE’s spatial community predictions.

**Location:** New world tropical and temperate plant communities.

**Methods:** We analytically derived and simulated METE’s spatially explicit expectations for the Sorensen index of community similarity. We then compared the distance decay of community similarity of 16 mapped plant communities to METE and the random placement model.

**Results:** The version of METE we examined was successful at capturing the general functional form of empirical distance decay relationships, a negative power function relationship between community similarity and distance. However, the semi-recursive approach consistently over-predicted the degree and rate of species turnover and yielded worse predictions than the random placement model.

**Main conclusions:** Our results suggest that while METE’s current spatial models accurately predict the spatial scaling of species occupancy, and therefore core ecological patterns like the species-area relationship, its semi-recursive form does not accurately characterize spatially-explicit patterns of correlation. More generally, this suggests that tests of spatial theories based only on the species-area relationship may appear to support the underlying theory despite significant deviations in important aspects of spatial structure.

## Introduction

Community structure can be characterized using a variety of macroecological relationships such as the species-abundance, body size, and species spatial distributions. Increasingly ecologists have recognized that many of these macroecological patterns are inter-related, and progress has been made toward unifying the predictions of multiple patterns using theoretical models (Storch *et al.*, 2008; McGill, 2010). One approach to predicting suites of macrecological patterns are process-based models such as niche and neutral dispersal models, which have the potential to provide biological insight into the process structuring ecological systems (Adler *et al.*, 2007). Alternatively, a new class of constraint-based models suggest that similar patterns may be produced by different sets of processes because the form of the predicted pattern is due to the existence of statistical constraints rather than directly reflecting detailed biological processes (Frank, 2009, 2014; McGill & Nekola, 2010; Locey & White, 2013).

The Maximum Entropy Theory of Ecology (METE) is a recent attempt to explain a number of ecological patterns from the statistical constraint perspective (Harte *et al.*, 2008, 2009; Harte, 2011; Harte & Newman, 2014). METE uses the principle of entropy maximization, that the most likely distribution is the one with the least information (i.e., the one closest to the uniform distribution) subject to a set of constraints (i.e., prior information), to predict distributions of species abundance, body size, and spatial structure. A frequentist perspective on the Maximum Entropy modeling approach is that every possible configuration of a system is equally likely; therefore, the probability of a particular distribution is directly proportional to the number of configurations that distribution is compatible with (Harte, 2011; Harte & Newman, 2014). The distribution with the largest number of compatible system configurations is the predicted most likely state of the system. In contrast to detailed biological models of community assembly, METE has no free parameters and only requires information on total community area, total number of individuals, total number of species, and total metabolic rate of all individuals to generate its predictions.

There is strong empirical support for METE’s predictions for the species abundance distribution and patterns related to the spatial distribution of individuals and species (Harte *et al.*, 2008, 2009; Harte, 2011; White *et al.*, 2012a; Xiao *et al.*, 2013; McGlinn *et al.*, 2013; Newman *et al.*, 2014). Specifically, METE has been successful at predicting spatially implicit patterns of community structure such as the species spatial abundance distribution and the species-area relationship (Harte *et al.*, 2008, 2009; McGlinn *et al.*, 2013). It has even been proposed that the METE spatial predictions yield a widely applicable universal species-area relationship (Harte *et al.*, 2009, 2013, but see Šizling *et al.*, 2011, 2013). However all of METE’s spatial predictions that have been tested focus on spatially implicit patterns that ignore spatial correlations. As a result the theory has not been evaluated using one of the core components of spatial structure. This is due in part to the fact that METE’s spatial correlation predictions have not been fully derived.

The most commonly studied ecological pattern that relies on these spatial correlations is the distance decay relationship (DDR) in which the similarity of species composition decreases with distance (Nekola & White, 1999). The DDR provides a spatially-explicit, community-level characterization of intra-specific aggregation patterns including correlations in space (Plotkin & Muller-Landau, 2002; Palmer, 2005; Morlon *et al.*, 2008; McGlinn & Palmer, 2011), and predicting the DDR is an important area of future development for METE because the DDR is necessary to accurately extrapolate community patterns to unsampled areas (Harte, 2011).

Here we explore METE’s spatially explicit predictions for the DDR by developing analytical and simulation based solutions and comparing them to empirical data. We build on the Hypothesis of Equal Allocation Probabilities (HEAP, Harte et al. 2005, Harte 2007) using an approach that combines elements of both a non-recursive and recursive version of METE (McGlinn et al. 2013). We test those predictions using data from 16 spatially explicit plant communities and compare METE’s performance to the classic Random Placement Model (RPM) in which individuals are randomly placed on the landscape (Coleman, 1981). Our approach provides a stronger evaluation of the performance of this model and whether it can explain patterns of spatial structure in the absence of detailed biological processes.

## Methods

METE has thus far been used to derive the probability that a random cell on a landscape will be occupied by a given number of individuals (i.e., the intra-specific spatial abundance distribution). Predictions for this distribution have been based either on recursively subdividing an area in half or on predicting species abundances directly at smaller scales (Harte, 2011; McGlinn *et al.*, 2013). In addition to the spatial abundance distribution, the DDR requires a prediction for the correlations in abundance among neighboring cells, which has proven difficult to derive for METE (Harte 2011).

### Developing METE’s Spatially Explicit Predictions

METE’s spatial predictions depend on two conditional probability distributions which are computed using independent applications of MaxEnt:

1. the species abundance distribution (SAD), Φ(*n* | *S*_0_, *N*_0_), the probability that a species has abundance *n* in a community with *S*_0_ species and *N*_0_ individuals, and
2. the intra-specific spatial abundance distribution, Π(*n* | *A*, *n*_0_, *A*_0_), the probability that *n* individuals of a species with *n*_0_ total individuals are located in a random quadrat of area *A* drawn from a total area *A*_0_.

The METE prediction for Φ is calculated using entropy maximization with constraints on the average number of individuals per species (*N*_0_/*S*_0_) and the maximum number of individuals *N*_0_ for a given species, which yields a truncated log-series abundance distribution (Harte *et al.*, 2008; Harte, 2011). The spatially implicit Π distribution is solved for using entropy maximization with constraints on the average number of individuals per unit area (*n*_0_/*A*_0_) and the maximum number of individuals *n*_0_ of a given species. Although METE requires information on total metabolic rate to derive its predictions, the exact value that this constraint takes has no influence on Φ and Π (Harte *et al.*, 2009; Harte, 2011).

Previous studies have downscaled (or upscaled) METE’s predictions using recursive and non-recursive approaches. Here we develop a spatially explicit approach to downscaling METE’s predictions that combines elements of both approaches and builds off an existing theoretical framework for modeling the DDR. With the recursive version of METE, Φ and Π are solved for at each successive halving or bisection of *A*_0_ until the area of interest is reached. After each bisection, Φ and Π are calculated and used to derive predicted values of average *S* and *N* at that scale which provide updated constraints for the next bisection (Harte et al. 2009).

Alternatively, a non-recursive approach can be used in which, Φ and Π at the spatial grain of interest can be solved for directly from the constraints placed at *A*_0_ (Harte et al. 2008). A semi-recursive approach is also possible in which Π is recursively downscaled but Φ is not. The semi-recursive predictions of METE have not been previously examined but this model builds directly on the existing theoretical derivations of the DDR by Harte (2007) for the Hypothesis of Equal Allocation Probabilities (HEAP). In Appendix A, Fig. A1 and A2 we examine how the semi-recursive formulation of METE differs from a previous examination of the METE recursive and non-recursive SARs (McGlinn et al. 2013), and in Appendix B we develop the analytical derivations of the semi-recursive formulation of the DDR.

In the semi-recursive formulation of the DDR, multi-cell correlations emerge from the spatially nested application of a recursive bisection scheme in which individuals are randomly placed in the left or right half of a cell at each bisection (Fig. 1). Biologically, this can be thought of as a sequentially dependent colonization rule in which individuals randomly choose to occupy the left or right side of an area depending on the existing number of individuals in each half (Harte et al. 2005, Harte 2007, and Conlisk *et al.* 2007). Our version of METE assumes that for a single bisection there is an equal likelihood for every possible spatial configuration of indistinguishable individuals (Eq. B1). Multi-cell spatial correlations emerge from this approach because the two cells that are formed from a common parent cell are adjacent to one another and are likely to be more similar in abundance than other cells on the landscape (Fig. 1). This approach has three important and inter-related limitations: 1) At each stage in the bisection algorithm, information about the cells surrounding the parent cell is ignored when determining allocations within the parent cell, 2) between-cell distance is defined in reference to an artificial bisection scheme which does not have a one-to-one correspondence with physical distance, and 3) the correlation between cells does not decrease smoothly with physical distance. Alternative approaches have been proposed for deriving the DDR for METE based on computing the single-cell Π distribution at two or more scales and then using the scaling of this marginal distribution to infer the probabilities of a given spatial configuration of abundance (Harte 2011). However, these approaches have yet to yield predictions for the DDR.

**Fig 1.**
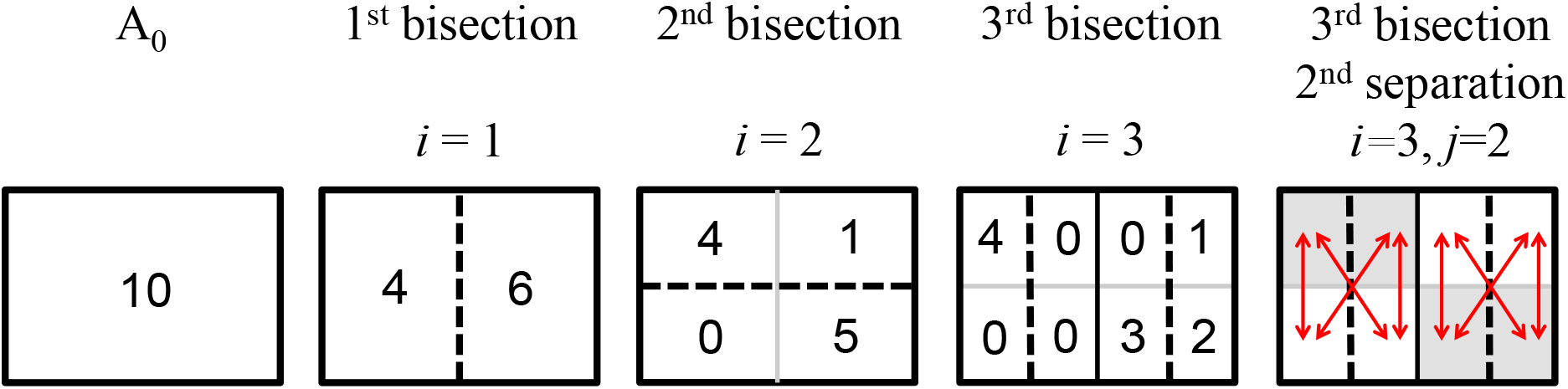
This diagram illustrates the “user rules” of how a landscape is bisected and how samples are compared for a given separation order. In this specific example, three bisections are used to generate a spatially explicit distribution of 10 individuals. In the last panel, the eight pairwise comparisons (arrows) at separation order of 2 for a scale of *A*_0_/2^3^ (i.e., *A*_*i*=3_, *D*_*j*=2_) are illustrated. When simulating random bisections the number of individuals distributed to the left or right of the bisection line is a random draw from a discrete uniform distribution.

The analytical forms of the semi-recursive formulation (Appendix B) are time-intensive to compute due to the multiple levels of recursion, ignore patterns of abundance (i.e., are formulated only in terms of presence-absence), and are not exact. An alternative approach to deriving semi-recursive METE predictions for the DDR is to use a spatially-explicit simulation.

### Spatially Explicit METE Simulation

To simulate semi-recursive METE’s spatial predictions, the equal probability rule (Eq. B1) that METE assumes when total area is halved is recursively applied starting at the anchor scale *A*_0_ and progressively bisecting the area until the finest spatial grain of interest is achieved (Fig. 1). Abundance in the simulation model can be parameterized using an observed SAD or using a random realization of the METE SAD given the values of *S*_0_ and *N*_0_. Once the abundances of the species are assigned, each species is independently spatially distributed. Because the equal probability rule requires that there is an equal probability of 0 to *n*_0_ individuals occurring on the left or right side of the total area *A*_0_, the number of individuals in the left side can be set as a draw from a discrete random uniform distribution between 0 and *n*_0_ and the remaining number of individuals are placed on the right hand side.

### Datasets

We used a database of 16 spatially explicit and contiguous community datasets compiled by McGlinn et al. (2013) to evaluate the DDR predictions of recursive METE (Table 1). All of the sites were terrestrial, woody plant communities with the exception of the serpentine grassland dataset which covered a terrestrial, herbaceous plant community. In the woody plant communities, all stems were recorded that were at least 10 mm in diameter at breast height (i.e., 1.4 m from the ground) with the exception of the Oosting and Cross Timbers sites where the minimum diameter was 20 and 25 mm respectively. Recursive METE only generates predictions for bisections of total area; therefore, we restricted our analysis to square or rectangular areas with a length-to-width ratio of 2:1. Two of the sites had irregular plot designs: Sherman and Cocoli. At these sites we partitioned the datasets into two 2:1 rectangles and analyzed each half independently and then averaged the results (see Supplemental Information: Fig. S1 in McGlinn et al. 2013). See McGlinn et al. (2013) for additional information on site selection criteria, and in particular their Supplemental Table 1, which provides a more complete description of the datasets used in our analysis.

**Table 1.** Summary of the habitat type and state variables of the vegetation datasets. The state variables are total area (*A*_0_), total abundance (*N*_0_) and total number of species (*S*_0_). *A*min and *A*max are the finest and coarsest areas (m^2^) examined. Data were collected on woody forest plants with the exception of the serpentine site which contained herbaceous grassland plants.

| Site name | Habitat type | Ref | $A_{\min}$ | $A_{\max}$ | $A_0$ | $N_0$ | $S_0$ |
| --- | --- | --- | --- | --- | --- | --- | --- |
| BCI | tropical | <sup>1-3</sup> | 61.0 | 62500 | 500000 | 205096 | 301 |
| Sherman | tropical | <sup>4</sup> | 2.4 | 625 | 20000 | 7623 | 175 |
| Cocoli | tropical | <sup>4</sup> | 2.4 | 625 | 20000 | 4326 | 139 |
| Luquillo | tropical | <sup>5</sup> | 15.3 | 15625 | 125000 | 32320 | 124 |
| Bryan | oak-hickory | <sup>6-8</sup> | 2.1 | 535 | 17113 | 3394 | 48 |
| Big Oak | oak-hickory | <sup>6-8</sup> | 2.4 | 625 | 20000 | 5469 | 40 |
| Oosting | oak-hickory | <sup>9</sup> | 16 | 4096 | 65536 | 8892 | 39 |
| Rocky | oak-hickory | <sup>6-8</sup> | 3.5 | 900 | 14400 | 3383 | 37 |
| Bormann | oak-hickory | <sup>6-8</sup> | 4.8 | 1225 | 19600 | 3879 | 30 |
| Wood Bridge | oak-hickory | <sup>6-8</sup> | 1.2 | 315 | 5041 | 758 | 19 |
| Bald Mtn. | oak-hickory | <sup>6-8</sup> | 2.4 | 156 | 5000 | 669 | 17 |
| Landsend | old field, pine | <sup>6-8</sup> | 1.0 | 264 | 8450 | 2139 | 41 |
| Graveyard | old field, pine | <sup>6-8</sup> | 2.4 | 625 | 10000 | 2584 | 36 |
| UCSC | mixed-evergreen | <sup>10</sup> | 5.4 | 1406 | 45000 | 5885 | 31 |
| Serpentine | serpentine | <sup>11</sup> | 0.3 | 4 | 64 | 37182 | 24 |
| Cross Timbers | oak woodland | <sup>12</sup> | 9.8 | 2500 | 40000 | 7625 | 7 |
| **Ranges** |  |  | 0.3-61.0 | 4-62500 | 64-500000 | 669-205096 | 7-301 |

<sup>1.</sup> Condit (1998), <sup>2.</sup> Hubbell et al. (1999), <sup>3.</sup> Hubbell et al. (2005), <sup>4.</sup> Condit et al. (2004), <sup>5.</sup> Zimmerman et al. (1994), <sup>6.</sup> Peet and Christensen (1987), <sup>7.</sup> McDonald et al. (2002), <sup>8.</sup> Xi et al. (2008), <sup>9.</sup> Palmer et al. (2007), <sup>10.</sup> Gilbert et al. (2010), <sup>11.</sup> Green et al. (2003), <sup>12.</sup> Arévalo (2013)

### Data Analysis

We compared the fit of METE with and without the observed SAD and the random placement model (RPM) to the empirical DDRs. The METE predictions represented averages of the abundance-based Sørensen index across 200 simulated communities. The abundance-based RPM predictions were generated by distributing the observed number of individuals of each species randomly in space and then computing the average abundance-based Sørensen index across 500 permutations (Morlon *et al.*, 2008).

The DDR is sensitive to the choice of the spatial grain of comparison (Nekola & White, 1999); so, we examined the DDR at several spatial grains for each dataset. We examined spatial grains resulting from 3-13 bisections of *A*_0_. To ensure that the samples at a given grain were square we only considered odd numbers of bisections when *A*_0_ was rectangular and even numbers of bisections when *A*_0_ was square. To ensure the best possible comparison between the observed data and METE and to avoid detecting unusual spatial artefacts in the METE predicted patterns we employed the “user rules” of Ostling et al. (2004) such that samples at a specific grain (i.e., level of bisection) were only compared if they were separated by a specific line of bisection (i.e., a given separation order, Fig. 1 and Appendix A, Fig. A3). This approach was taken rather than the standard method of constructing the DDR from all possible pairwise sample comparisons without reference to an imposed bisection scheme. We computed geographic distance by averaging the distance between all the compared samples compared at a given separation order. For the Crosstimbers study site we were not able to examine the DDR based on the METE SAD because of difficulty in generating random realizations of the METE SAD needed for the community simulator when *S*_0_ is less than approximately 10. Typically averages of community similarity are used to examine the geometry of the DDR; however, in some cases the distribution of the similarity metric may be strongly skewed and therefore we computed both averages and medians of community similarity at each separation order.

We used weighted least squares (WLS) regression to account for differences in the number of pairwise comparisons at different spatial lags (there are many more comparisons at short lags) when fitting the power and exponential models of the DDR (Venables & Ripley, 2002). We examined the power model and exponential models because they are the simplest statistical models of the DDR, and it was recently suggested that at fine spatial scales the DDR should be best approximated by a power model (Nekola & White, 1999; Nekola & McGill, 2014).

We checked that our results were consistent with the results provided in previous studies (Harte, 2007, Fig. 6.7 and 6.8, 2011, Fig. 4.1), and that the DDR generated by the community simulator closely agreed with the analytical solution Eq. B5 (Appendix B, Fig. B1). The code to recreate the analysis is provided as Appendix D and at the following publicly available repository: http://dx.doi.org/10.6084/m9.figshare.978918.

## Results

In general, the semi-recursive METE distance decay relationship (DDR) provided a poor fit to the empirical DDR (Figs. 2 and 3). The average and median community similarity results were highly correlated (*r* = 0.98) and generated qualitatively similar results (Appendix A Figs. A5 and A8); therefore, we focus on the results based on averaging similarity. While the METE DDRs exhibited the general functional form of the empirical DDRs, an approximately power-law decrease in similarity with distance, they typically had lower intercepts and steeper slopes than the empirical DDRs (Fig. 2, Appendix A, Fig. A4 and A6). Both the empirical and METE predicted DDR were better approximated by power rather than exponential models (Appendix A, Fig. A6). METE converged towards reasonable predictions at fine spatial grains; however, this is to be expected because at these scales similarity in both the observed and predicted patterns must converge to zero due to low individual density (grey points in Fig. 3A,B). This is because when individual density is low the probability of samples sharing species decreases rapidly simply due to chance. The RPM is known to be a poor model for distance decay because it does not exhibit a decrease in similarity with distance. However, it fit the empirical DDR slightly better than METE (Figs. 2 and 3).

**Fig 2.**
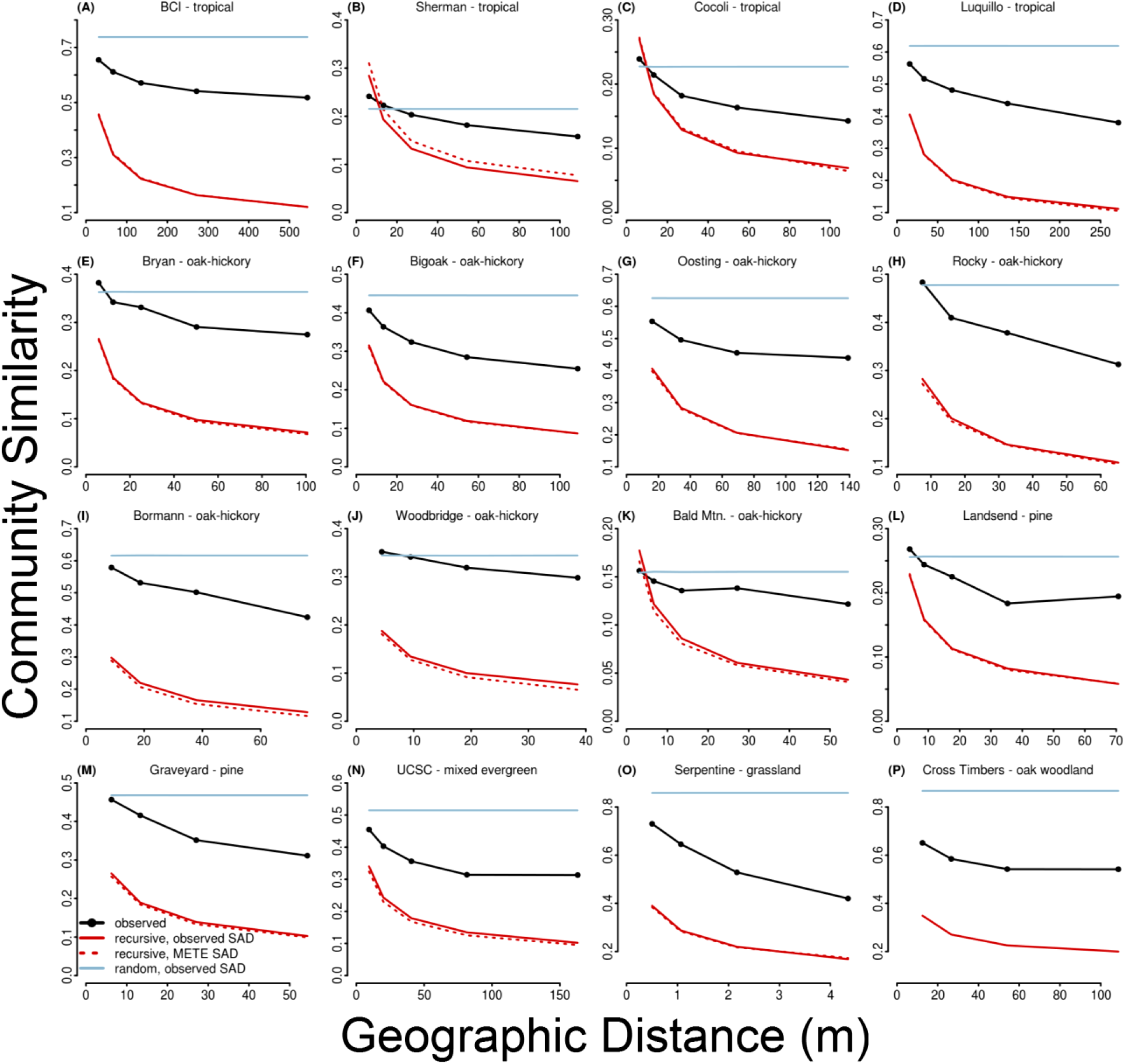
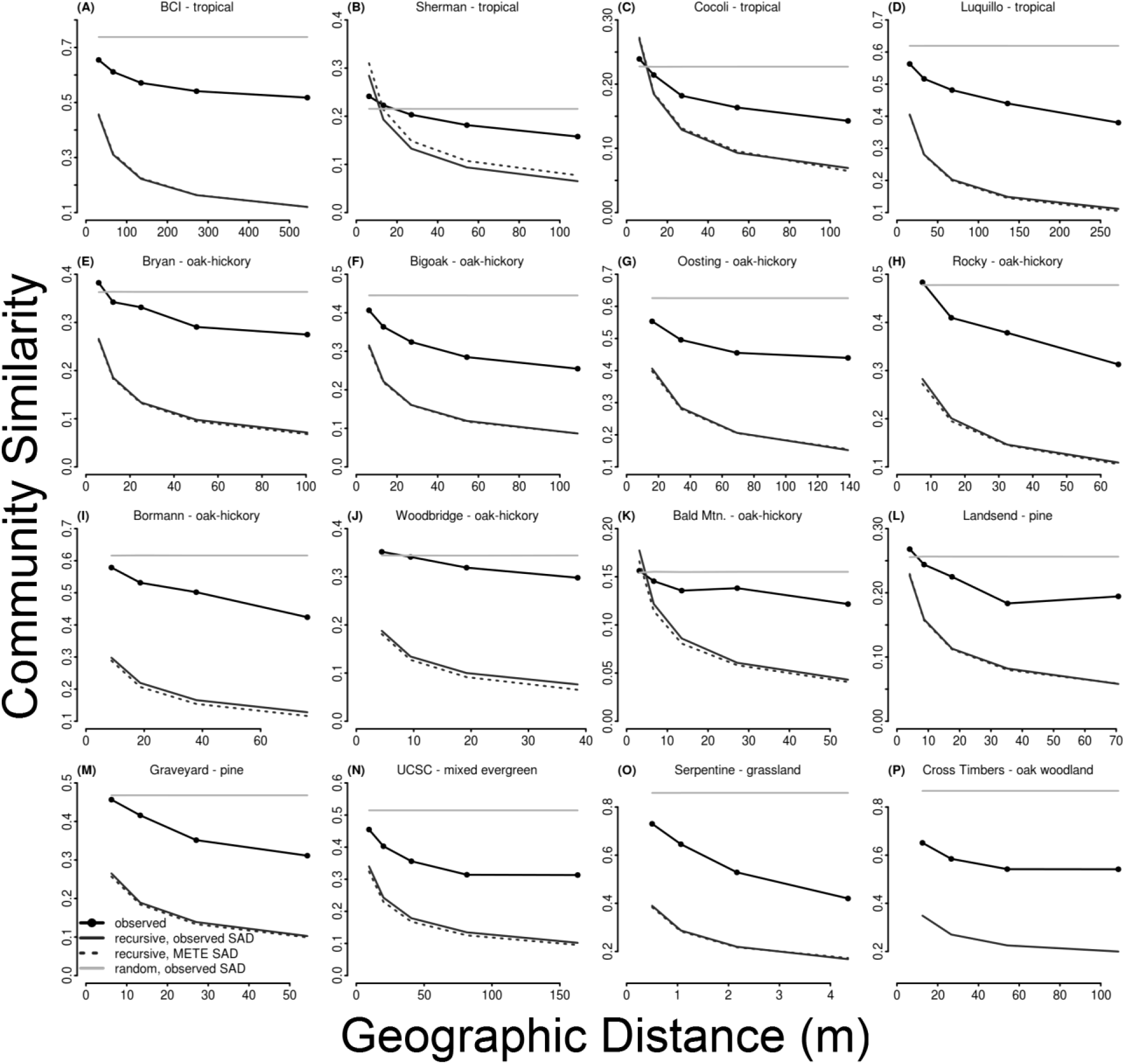
The observed (black line with dots) and predicted distance decay relationships (METE: dark grey lines, solid for the observed SAD, dashed for the METE SAD; random placement: light grey line) for each site at a single spatial grain. Community similarity represents the average of the abundance-based Sørensen index for each spatial lag. The spatial grain displayed was taken at either 8 or 9 bisections of the total area depending on whether the total extent was a square or a rectangle respectively. Geographic distance was calculated as the average physical distance between the samples compared at given separation order (see Methods and Fig. 1 for additional information).

**Fig 3.**
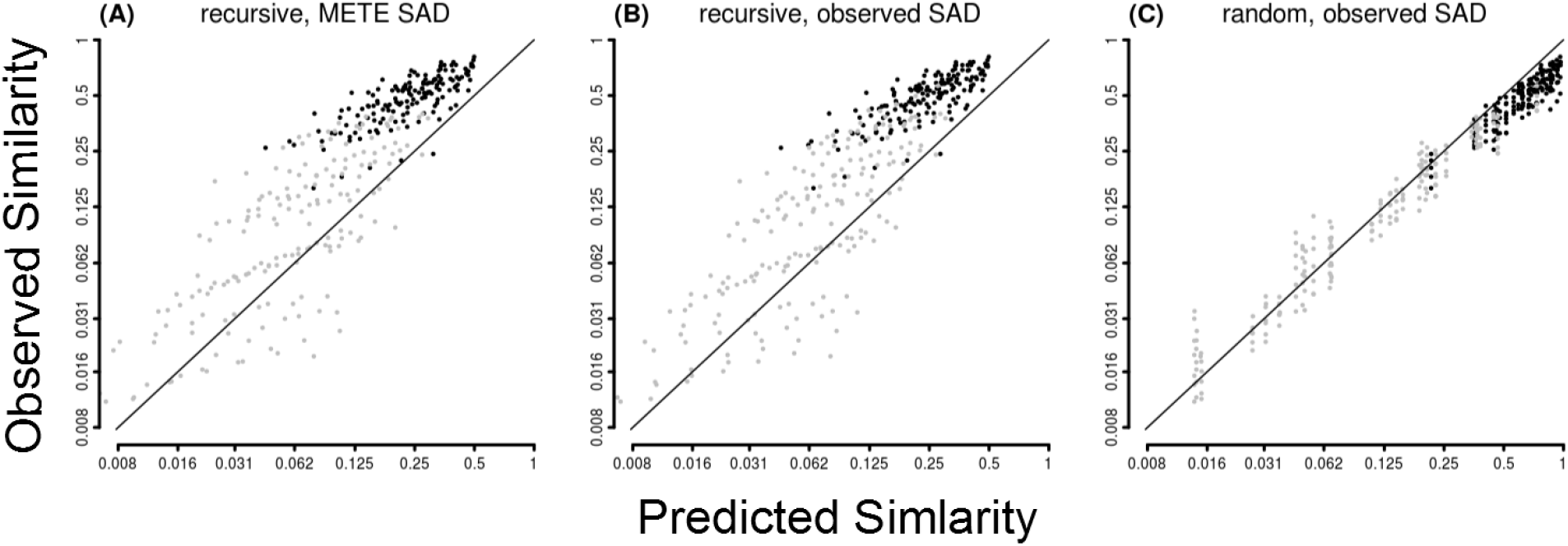
The log-log transformed one-to-one plots of the predicted and observed abundance-based Sørensen similarity values for the three models across all distances and spatial grains. The solid line is the one-to-one line. The grey points represent values from spatial grains in which the average individual density was low (i.e., less than 10 individuals) and thus both the observed and predicted similarities must be close to zero simply because of a sampling effect.

The METE DDR was not strongly influenced by the choice of using the observed or the METE SAD (Figs. 2 and 3A,B). The METE SAD typically yielded a DDR with a slightly lower intercept with the exception of the four tropical sites where it produced DDRs with slightly higher intercepts. In general, we did not observe strong consistent differences between the habitat types (Fig. 2, Appendix A, Fig. A7).

Our formulation of a semi-recursive METE produced SARs that generally agreed (i.e., within the 95% CI) with the recursive and non-recursive formulations of METE (Harte et al. 2009); however, it did appear that the semi-recursive approach systematically deviated towards lower richness at fine spatial scales which is consistent with predicting stronger patterns of spatial aggregation compared to the other formulations of METE (Appendix A, Fig. A1 and A2).

## Discussion

The semi-recursive METE distance decay relationship (DDR) was well approximated by a decreasing power function, and thus consistent with the general form of empirical DDRs, but it provided a poor fit to empirical data. Specifically, the slope and the intercept of this power function deviate substantially from empirical data resulting in a poor fit. These deviations contrast with a number of studies showing that the theory successfully predicts both the Π distribution and the SAR (Harte *et al.*, 2008, 2009; Harte, 2011; McGlinn *et al.*, 2013; but see Šizling *et al.*, 2011). Both Π and the SAR are influenced by the spatially explicit pattern of intraspecific aggregation but neither pattern reflects inter-quadrat correlations and therefore they represent coarse metrics of spatial structure. The combination of a well fit SAR and a poorly fit DDR suggests that the current version of METE accurately characterizes average occupancy, but fails to characterize the spatial relationships among cells (McGeoch & Gaston, 2002; Storch *et al.*, 2003; McGlinn & Hurlbert, 2012; Nekola & McGill, 2014).

These results only apply directly to the particular HEAP-based semi-recursive version of the spatial METE theory, which represents a middle ground in terms of approach between Harte et al. (2008) and Harte *et al.* (2009). Other approaches to deriving the METE DDR may perform better than the semi-recursive approach if they can be developed. It has been suggested that there is no *a priori* reason to prefer one version of the theory and that the best way to choose among the different versions is empirically (Haegeman & Etienne, 2010; Harte, 2011). However, the traditionally defined recursive and non-recursive versions of METE have shortcomings with respect to how their assumptions and predictions are scaled, and the semi-recursive approach we defined is limited by its dependence on an artificial bisection scheme. Specifically the recursive approach predicts that the SAD has the same functional form, a truncated log-series, at all scales. This is problematic because SADs are typically not scale-invariant if, as METE predicts, species display intraspecific spatial aggregation (Green & Plotkin, 2007; Šizling *et al.*, 2009). The non-recursive approach does not suffer from this problem because the SAD is only solved for at the anchor scale; however, Haegemann and Etienne (2010) found that the non-recursive predictions for a multi-cell generalization of the Π distribution were scale-inconsistent. The semi-recursive approach does not suffer from this shortcoming because its multi-cell form (see Eq. 2.2 in Conlisk *et al.*, 2007) is only defined over the set of bisections that are consistent with a landscape in which *n*_0_ individuals are distributed (see Appendix C for proof). However, the set of bisections is artificial and multi-cell correlations only emerge from this approach in reference to bisection distance rather than directly to physical distance between cells such that cells have equal magnitude of correlation regardless of their physical distance if they have equivalent separation orders (see Conlisk *et al.*, 2007 for a critique of distances defined by separation indices). An important future direction for METE is to attempt to develop spatial multi-cell predictions using approaches that avoid these shortcomings and the two approaches suggested by Harte (2011) for deriving the METE DDR may provide a useful starting point for future development.

Our results suggest that semi-recursive METE differs from spatial patterns observed in nature. This deviation could indicate that the emergent statistical approach to modeling spatial structure is incorrect, with specific biological processes such as dispersal limitation or environmental filtering directly controlling spatial correlation (Condit *et al.*, 2002; Gilbert & Lechowicz, 2004; Karst *et al.*, 2005; Seidler & Plotkin, 2006; Chase, 2007; McGlinn & Palmer, 2011). Alternatively it could mean that while the general idea underlying the theory is valid, the specific formulation is wrong. For example it could be that the approaches outlined by Harte (2011) that are more sophisticated in how they handle spatial correlations will be more appropriate or that a generalized version of this kind of recursive approach like that developed by Conlisk et al. (2007) in which the degree of aggregation is a tunable parameter will capture the reality of biological systems more precisely. However, process-, and constraint-based models should not necessarily be treated as mutually exclusive. For example other process-based theories make power-law like predictions for the form of the DDR. In fact, it has recently been suggested that at fine spatial scales most theories will make predictions that are approximately power-law in nature (Nekola & McGill, 2014). This means that simply noting power-law like DDR relationships does not provide a strong method for differentiating among theories. In fact, had we simply looked for power-law like behavior we would have concluded that the semi-recursive METE was consistent with empirical data. However, one of the properties that makes METE such a strong theory is that it makes specific predictions for precise parameters as well as general forms of empirical relationships. This allows it to be more rigorously compared to data and to other theories that predict different parameters values for a similar general form of the DDR (e.g., neutral theory).

Our results mirror those of Xiao et al. (2013) and Newman et al. (2014) evaluating the non-spatial aspects of METE. All three studies show that when evaluating the theory using multiple patterns simultaneously some of the predictions perform well and some perform poorly. It is inherently difficult for theories to predict large numbers of patterns simultaneously, which is why evaluating theory in this way provides stronger tests than evaluating single patterns (McGill, 2003; McGill *et al.*, 2006). General theories like METE that make multiple predictions are therefore both easier to evaluate and also more broadly useful since they allow a large number of patterns to be predicted from a relatively small amount of information. Because there are many patterns to evaluate it is also more likely that deviations from theory will be identified (White *et al.*, 2012). In some cases these deviations may indicate that the theory is fundamentally unsound, but in others it may suggest modifications to the theory to address the observed deviations (White *et al.*, 2012). Whether METE can be modified to address the observed deviations from empirical data remains to be seen. In the case of the DDR, despite its generality, there are a limited number of models that attempt to predict the DDR from first principles (Chave & Leigh, 2002; Condit *et al.*, 2002; Zillio *et al.*, 2005; Harte, 2007, 2011; Nekola & McGill, 2014), which means that it may be worth pursuing the METE approach further.

METE is one of several general theories in ecology that make many predictions for many aspects of ecological community structure based on only a small amount of information. Our analysis of the semi-recursive formulation of METE’s spatially explicit prediction for the DDR suggests that this form of the theory over-predicts the strength of spatial correlation. These results coupled with studies of the species-area relationship suggest that semi-recursive METE accurately predicts the scaling of species occupancy but not spatial correlation. More generally, our results demonstrate that tests of spatial theories that focus solely on the species-area relationship and related patterns are only evaluating part of the spatial pattern, the distribution of occupancy among cells. Evaluating these theories using the DDR in addition to the SAR will help identify cases where the theories are correctly identifying some aspects of spatial structure, but not others, and thus yield stronger tests of the underlying theory. In some cases this will require extending the theory to make additional predictions, but this effort will provide both more testable and more usable theories.

## Acknowledgements

We thank John Harte for extensive conversations on the use of maximum entropy in ecology. Robert K. Peet provided data for the oak-hickory and old field pine forests. Jessica Green provided data for the serpentine grassland data. José Arévalo provided data for the oak woodland data. The BCI forest dynamics research project was made possible by National Science Foundation grants to Stephen P. Hubbell: DEB-0640386, DEB-0425651,DEB-0346488, DEB-0129874, DEB-00753102, DEB-9909347, DEB-9615226, DEB-9615226, DEB-9405933, DEB-9221033, DEB-9100058, DEB-8906869, DEB-8605042,DEB-8206992, DEB-7922197, support from the Center for Tropical Forest Science, the Smithsonian Tropical Research Institute, the John D. and Catherine T. MacArthur Foundation, the Mellon Foundation, the Small World Institute Fund, and numerous private individuals, and through the hard work of over 100 people from 10 countries over the past two decades. The plot project is part the Center for Tropical Forest Science, a global network of large-scale demographic tree plots. The Luquillo Experimental Forest Long-Term Ecological Research Program, supported by the U.S. National Science Foundation, the University of Puerto Rico, and the International Institute of Tropical Forestry. This research was supported by a CAREER grant from the U.S. National Science Foundation to EPW (DEB-0953694).

## Biosketch

Weecology is founded on the belief that better communication and collaboration between empirical and quantitative scientists is necessary for tackling many of the big scientific challenges in ecology. The purpose of Weecology is to facilitate collaborative research through a variety of mechanisms including shared resources, expertise, and web-based collaborative tools. Our hope is help scientists (and ourselves) collaborate and communicate across disciplinary divides and generate higher-quality novel research as a result.

